# Cryo-EM structure of a type VI secretion system delivered membrane-depolarizing toxin involved in bacterial antagonism

**DOI:** 10.1101/2025.04.30.651342

**Authors:** Jake Colautti, Dmitry Shatskiy, Eileen Bates, Nathan P. Bullen, Alexander Belyy, John C. Whitney

**Affiliations:** Michael DeGroote Institute for Infectious Disease Research, McMaster University, Hamilton, ON, L8S 4K1, Canada; Department of Biochemistry and Biomedical Sciences, McMaster University, Hamilton, ON, L8S 4K1, Canada; Membrane Enzymology Group, Groningen Institute of Biomolecular Sciences and Biotechnology (GBB), Faculty of Science and Engineering, University of Groningen, Groningen, The Netherlands; David Braley Center for Antibiotic Discovery, McMaster University, Hamilton, ON, L8S 4K1, Canada

**Keywords:** *Pseudomonas aeruginosa*, type VI secretion systems, membrane-depolarizing toxins, cryo-EM

## Abstract

Many Gram-negative bacteria use type VI secretion systems (T6SSs) to deliver toxic effector proteins into neighboring competitor cells. Members of the VasX protein family, such as VasX from *Vibrio cholerae* and Tke5 from *Pseudomonas putida*, disrupt the inner membrane of target cells by forming ion-permeable channels that dissipate the proton motive force, thereby interfering with essential physiological processes. However, the molecular structure of any VasX family effector has remained unknown. Here, we present a cryo-EM structure of Ptx2, a recently identified VasX family effector exported by a T6SS of *Pseudomonas aeruginosa*. Our structure reveals that Ptx2 is an elongated, multi-domain protein that bears little resemblance to proteins of known function. Notably, the apparent flexibility of its domains suggests that Ptx2, like other membrane-depolarizing toxins, undergoes substantial conformational changes to facilitate membrane insertion. Guided by these predicted structural rearrangements, we used mutagenesis coupled with phenotypic assays to identify key features required for its toxic activity. Together, these findings provide the first molecular level insights into the structure and mechanism of VasX family effectors and expand our understanding of how these proteins contribute to interbacterial antagonism.

## Introduction

To survive in densely populated environments where competition for resources is intense, many bacteria secrete toxic proteins into competitor cells to gain a fitness advantage (Klein et al., 2020; Ruhe et al., 2020). In Gram-negative bacteria, contact-dependent toxin delivery is frequently facilitated by type VI secretion systems (T6SSs), which are contractile protein export apparatuses that span the diderm cell envelope. Upon contraction, T6SSs inject a needle-like protein complex decorated with toxic effector proteins directly into adjacent competitor cells (Cherrak et al., 2019; Silverman et al., 2012). Following their injection, these effectors disrupt essential facets of cellular physiology, including cell division, central metabolism, or cell envelope integrity, leading to target cell death (Russell et al., 2014). To protect against self-killing or killing of genetically identical kin cells, T6SS effectors are invariably encoded alongside cognate immunity proteins that bind to and inactivate their associated toxins. Thus, T6SSs enable bacteria to selectively inhibit the growth of nearby competitor organisms and gain access to limited resources in their environmental niche (Chatzidaki-Livanis et al., 2016; Sheahan et al., 2024; Speare et al., 2018; Wexler et al., 2016).

Although many T6SS effectors act as enzymes that degrade or modify essential cellular substrates, several effector families have been identified that do not display enzymatic activity but instead act by depolarizing the bacterial inner membrane, thus dissipating the proton motive force (PMF) (Gonzalez-Magana et al., 2022; LaCourse et al., 2018; Mariano et al., 2019; Miyata et al., 2013; Velázquez et al., 2025). This ubiquitous ion gradient provides the energy required for several important cellular activities, including ATP synthesis and nutrient uptake, and is therefore essential for bacterial survival (Mitchell, 1966). The PMF comprises two components: an electrical potential, referred to as Δψ, and a proton gradient, termed ΔpH. By acting as selective metal ion or proton channels, membrane-depolarizing toxins dissipate Δψ or ΔpH, respectively, arresting processes that rely on the PMF (Ulhuq and Mariano, 2022). Although several families of membrane-depolarizing effectors have been found to transit the T6SS and other antibacterial protein secretion systems, to date, the structures of these proteins and the mechanisms by which they conduct ions across bacterial membranes remain elusive.

Unlike effectors exported by T6SSs, membrane-depolarizing toxin families associated with other bacterial competition systems or that function as host cell-targeting virulence factors have been the focus of decades of mechanistic study, which has revealed several general principles that govern their activity (Dal Peraro and van der Goot, 2016; Ulhuq and Mariano, 2022). For example, most membrane-depolarizing toxins adopt two distinct conformations: a soluble conformation that enables these proteins to be secreted through an aqueous environment, and a membrane-inserted conformation that represents the active state of the toxin. In many instances, membrane-depolarizing toxins oligomerize upon inserting into target cell membranes, thus forming ion-conducting channels in which each protomer contributes a single secondary structure motif to the overall oligomer (Mueller et al., 2009; Song et al., 1996; Tanaka et al., 2015). Often, only a small region of a membrane-depolarizing toxin inserts into the target cell membrane and contributes to the ion channel; the remainder of the protein serves a chaperone-like function, stabilizing the membrane-inserting region prior to insertion. However, the structural studies that support this model have primarily focused on membrane-depolarizing bacterial virulence factors and diffusible antibacterial proteins known as colicins, both of which act in a cell contact-independent manner (Cascales et al., 2007). A mechanistic understanding of how membrane-depolarizing T6SS effectors insert into bacterial membranes and conduct ions across these membranes is therefore lacking.

We recently discovered Ptx2, an antibacterial T6SS effector secreted by the opportunistic human pathogen *Pseudomonas aeruginosa* (Colautti et al., 2024). Although its mechanism of action has not been demonstrated, Ptx2 is homologous to proteins belonging to the VasX family of membrane-depolarizing T6SS effectors, which includes the previously characterized *Vibrio cholerae* effector VasX and the *Pseudomonas putida* effector Tke5 (Miyata et al., 2011; Miyata *et al*., 2013; Russell et al., 2012; Velázquez *et al*., 2025). Both VasX and Tke5 display potent antibacterial activity when delivered to the periplasm of susceptible cells, whereupon they insert into bacterial membranes and dissipate the PMF. Tke5 has recently been found to function as an ion channel that selectively permeabilizes the bacterial inner membrane to both cations and anions, disrupting the PMF by dissipating Δψ. However, despite over a decade of research on antibacterial T6SS effectors belonging to this protein family, the molecular structure of VasX family proteins remains unknown.

Here, we present a cryogenic electron microscopy (cryo-EM) structure of Ptx2 in the soluble state. This structure reveals that Ptx2 adopts an elongated disk-shaped architecture comprised of multiple domains that bear little resemblance to proteins of known function. We further demonstrate that like other VasX family proteins, Ptx2 functions as an ion channel that disrupts Δψ, thus dissipating the proton motive force and inhibiting competitor cell growth. Interestingly, AlphaFold3 confidently predicts an alternative conformation of Ptx2 with a solvent exposed hydrophobic transmembrane region, potentially illustrating an ion-conducting pore state. By comparing our experimental structure of Ptx2 in its soluble conformation to the predicted structure in its membrane-inserted state, we identify key features of the toxin that are likely responsible for conformational changes between these two states. Finally, we demonstrate that Ptx2 is recruited to the H2-T6SS through its N-terminal MIX domain, providing a mechanistic explanation for how MIX domains recruit effectors to the T6SS apparatus. Together, our findings shed light on the molecular mechanisms by which the VasX family of proteins are recruited to the T6SS apparatus and how they dissipate the PMF to inhibit growth upon delivery into susceptible bacteria.

## Results

### Ptx2 acts from the bacterial periplasm

*P. aeruginosa* encodes three evolutionarily distinct T6SSs, referred to as the H1-, H2-, and H3-T6SS, which each export a unique effector repertoire and are therefore thought to function in distinct biological contexts (Barret et al., 2011; Colautti et al., 2025; Mougous et al., 2006). We recently discovered Ptx2, an antibacterial T6SS effector secreted by the H2-T6SS of the hypervirulent *P. aeruginosa* strain PA14, and Pti2, its cognate immunity protein (Colautti *et al*., 2024; Lee et al., 2006). However, the mechanism of action of this effector remains unknown. To identify homologs of Ptx2 that might provide clues to its function, we queried the sequence of this protein against the non-redundant protein database using an iterative hidden-Markov model approach (Zimmermann et al., 2018). This search revealed that Ptx2 is distantly related to the *Vibrio cholerae* T6SS effector VasX and the *Pseudomonas putida* effector Tke5 (17.3% sequence identity to VasX, 19.0% sequence identity to Tke5; supplementary file S1). VasX was among the first identified T6SS effectors and acts as an ion channel that dissipates the PMF, thus killing both bacterial and eukaryotic cells (Miyata *et al*., 2011; Miyata *et al*., 2013). Tke5 has recently been found to act by a similar mechanism and was further shown to form multi-ionic pores in isolated lipid bilayers that display some preference for cations over anions (Velázquez *et al*., 2025). Additionally, a VasX family effector has been shown to transit the bacteria-targeting T6SS-1 of *Burkholderia thailandensis*, although the function of this effector has yet to be investigated (Russell *et al*., 2012). Together, these proteins represent the characterized members of the VasX protein family, a widespread group of T6SS effectors that likely share a membrane-depolarizing mechanism of action (Velázquez *et al*., 2025). The sequence homology between Ptx2 and these VasX family proteins suggested to us that Ptx2 also acts as a membrane-depolarizing toxin.

Like other membrane-depolarizing T6SS effectors, VasX and Tke5 both act from the bacterial periplasm. If Ptx2 indeed acts by a similar mechanism, we would expect it to also exert its toxic effects from this cellular compartment. We previously discovered that in contrast to most described T6SS effectors, milligram quantities of Ptx2 can be expressed and purified from *E. coli*, suggesting that Ptx2 is not toxic in the bacterial cytoplasm (Colautti *et al*., 2024). To test the hypothesis that Ptx2 instead acts from the periplasm, we fused a twin-<u>a</u>rginine translocation (Tat) signal peptide to Ptx2 to target it to the periplasm of *E. coli*. Consistent with our previous findings, expression of native cytoplasmic Ptx2 (Ptx2_cyto_) did not impair the growth of *E. coli* (Figure 1B). By contrast, ectopic delivery of Ptx2 in the periplasm (Ptx2_peri_) potently inhibits bacterial growth. This observation suggests that like other VasX family proteins, Ptx2 acts from the periplasm.

**Figure 1:**
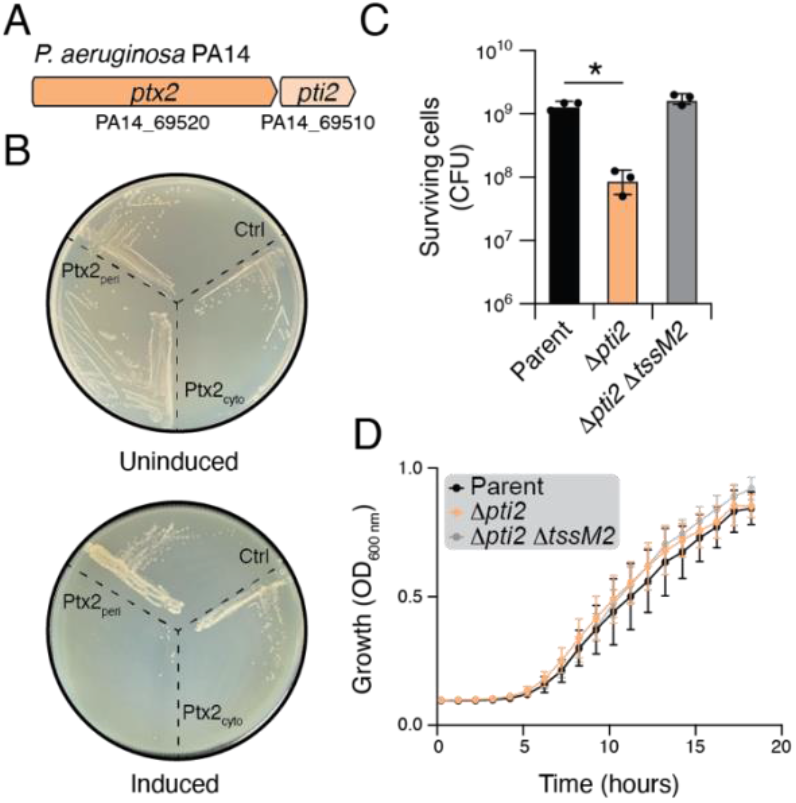
Ptx2 exerts its antibacterial activity from the periplasm. A) Schematic representation of the genes encoding Ptx2 and its cognate immunity protein Pti2. B) *E. coli* strains harbouring the indicated expression vectors grown on solid media in either inducing or non-inducing conditions. C) Viability of the indicated *P. aeruginosa* strains after 20 hours of growth on solid media. Mean colony forming units (CFUs)/mL were compared using an ordinary one-way ANOVA with multiple comparisons to the parent strain. D) Growth of the indicated *P. aeruginosa* strain in liquid media. In C and D, parent represents *P. aeruginosa* PA14 *ΔrsmA ΔamrZ*, error bars represent mean +/-SEM, n=3 biological replicates.

To further explore our hypothesis that Ptx2 is a periplasmic toxin, we next sought to assess if a *P. aeruginosa* strain lacking the Ptx2-specific immunity gene, *pti2*, undergoes T6SS-mediated intercellular self-intoxication. Because T6SS effectors are loaded onto the T6SS apparatus in the bacterial cytoplasm and are delivered directly into target cells in a single step event that bypasses the periplasm, effectors that act in the periplasm never access this cellular compartment in the producing cell (Russell et al., 2011). Immunity proteins that protect against such effectors are therefore non-essential for survival in conditions that are not conducive to T6SS-mediated protein delivery, such as growth in liquid media. Consequently, if Ptx2 indeed acts in the periplasm, we would expect a Δ*pti2* strain to survive in liquid culture but exhibit impaired growth on solid media where contacting cells would deliver Ptx2 into one another via their H2-T6SSs. We therefore constructed a *P. aeruginosa* Δ*pti2* strain and evaluated its growth under conditions that promote or impair T6SS-mediated protein delivery. For these experiments, we employed a genetic background lacking the H2-T6SS repressors *rsmA* and *amrZ*, which had previously been reported to express the H2-T6SS genes including the *ptx2-pti2* bicistron (Allsopp et al., 2017; Colautti *et al*., 2024). As predicted, we found that this strain displays impaired growth on solid media but grows comparably to its isogenic parent strain in liquid culture (Figure 1D, E). Furthermore, genetic inactivation of the H2-T6SS through deletion of the structural gene *tssM2* alleviates this growth defect, confirming that the observed growth inhibition arises from intercellular delivery of Ptx2 by the H2-T6SS. Because this property is a unique feature of T6SS effectors that act from the periplasm, we conclude that Ptx2 exerts its toxic activity from this cellular compartment.

### Ptx2 disrupts the electrical component of the proton motive force

Having established that Ptx2 acts in the bacterial periplasm, we next sought to determine if it disrupts the PMF of susceptible cells during T6SS-mediated bacterial competition. Previously characterized VasX family effectors act as membrane channels that facilitate the influx of ions from the extracellular environment into the cytoplasm, thus disrupting the electrical component of the proton motive force, Δψ. Tke5 was recently shown to function as a multi-ionic channel that facilitates the passive diffusion of both cations and to a lesser extent, anions, across isolated lipid bilayers (Velázquez *et al*., 2025). Because sodium ions are actively extruded from bacterial cells, we hypothesized that the toxicity of Ptx2 may partly arise from the passive transport of sodium ions from the extracellular environment into the cytoplasm, which would depolarize the cytoplasmic membrane and thus disrupt the PMF. In this model, we would expect Ptx2 toxicity to be ameliorated in the absence of extracellular sodium because the sodium concentration gradient between the extracellular environment and the cytoplasm would be diminished under these conditions and passive sodium influx would thus be reduced. To test this hypothesis, we conducted interbacterial competition experiments between a donor strain that constitutively expresses H2-T6SS genes and an isogenic Ptx2-susceptible recipient strain (*Δptx2Δpti2*) in the presence or absence of extracellular sodium. Consistent with our hypothesis, the Ptx2-dependent competitive fitness of the donor strain against the recipient strain is abrogated in media lacking extracellular monovalent cations that instead contains sucrose as an osmotic protectant (Figure 2A). Furthermore, the growth defect observed in our self-intoxicating Δ*pti2* strain is suppressed in the absence of sodium ions in the growth media (Figure 2B). These observations are consistent with the established role of VasX family effectors as ion channels that selectively disrupt the electrical component of the PMF (Miyata *et al*., 2013; Velázquez *et al*., 2025). Additionally, we conducted these competition experiments in media containing potassium rather than sodium. The extrusion of sodium into the environment is coupled with the cytoplasmic uptake of potassium. Therefore, membrane-depolarizing toxins that predominantly act by permitting passive efflux of cytoplasmic potassium are impaired when potassium is supplemented in the growth medium (LaCourse *et al*., 2018). Interestingly, the Ptx2-dependent fitness advantage of the donor strain is retained in the presence of extracellular KCl as is the growth defect of our Δ*pti2* strain (Figure 2A, B). This observation suggests that the efflux of cytoplasmic potassium is not a major contributor to Ptx2 toxicity during bacterial competition. Because VasX family proteins are known to conduct anions, albeit to a lesser extent than cations, it is possible that the toxicity of Ptx2 under these conditions arises partly from the passive transport of chloride ions into the cytoplasm where the concentration of this anion is typically low (Schultz et al., 1962). Together, these observations suggest that like other VasX-family proteins, Ptx2 acts as an ion channel to disrupt Δψ during T6SS-mediated interbacterial competition.

**Figure 2:**
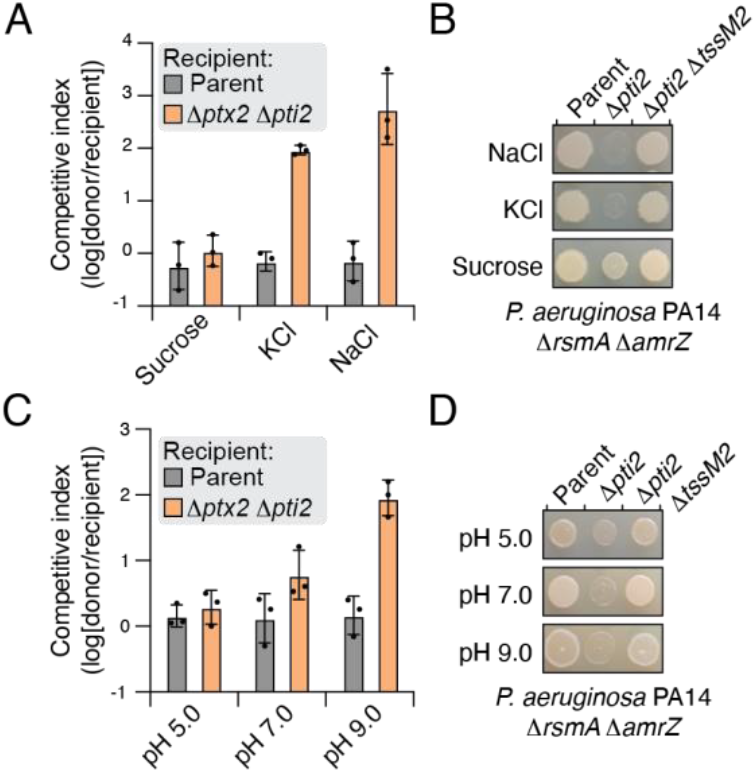
Ptx2 toxicity requires extracellular monovalent cations and is inhibited under acidic conditions. A, C) Outcome of intraspecific growth competitions between the parental donor strain and the indicated *P. aeruginosa* PA14 recipient strains. The competitive index is calculated as a change (final/initial) in the donor-to-recipient ratio. Error bars represent mean +/-SEM, n=3 biological replicates. Asterisks indicate statistically significant differences between the indicated donor and recipient strain (*p*<0.05). Differences between groups were calculated with a one-tailed homoscedastic T test comparing the competitive index of the parent vs the *Δptx2 Δpti2* strains. B) Growth of the indicated *P. aeruginosa* PA14 strains on solid media containing 150 mM NaCl, 150 mM KCl, or 300 mM sucrose. D). Growth of the indicated *P. aeruginosa* PA14 strains on solid media buffered with sodium phosphate pH 5.0, 7.0, or 9.0. In all panels, parent represents *P. aeruginosa* PA14 *ΔrsmA ΔamrZ*.

We next examined the role of extracellular pH on Ptx2-mediated toxicity. The chemical component of the PMF, ΔpH, is enhanced in an acidic environment and diminished under alkaline conditions (Booth, 1985). As a result, Δψ accounts for a greater proportion of the total PMF under alkaline conditions than under acidic conditions. Therefore, protein toxins and small-molecule antibiotics that act by selectively dissipating Δψ are potentiated under alkaline conditions, where Δψ contributes more strongly to the total PMF, and inhibited under acidic conditions (Farha et al., 2013). Consistent with the hypothesis that Ptx2 acts by dissipating Δψ, the Ptx2-dependent fitness advantage of the parental strain against the Ptx2-susceptible recipient is completely abrogated at pH 5.0 as is the growth defect of the self-intoxicating Δ*pti2* strain (Figure 2C, D). These observations lead us to conclude that T6SS-mediated delivery of Ptx2 selectively dissipates Δψ to disrupt the PMF. This finding is consistent with the recently established activity of VasX family proteins, which are known to form ion-permeable channels in the membrane that disrupt the PMF (Velázquez *et al*., 2025).

### Cryo-EM structure of soluble Ptx2

Although many families of membrane-depolarizing T6SS effectors have been identified, none have been structurally characterized (Gonzalez-Magana *et al*., 2022; LaCourse *et al*., 2018; Mariano *et al*., 2019; Miyata *et al*., 2013). To understand the molecular architecture of Ptx2, we heterologously expressed and purified this protein from *E. coli* and determined its structure using single-particle cryogenic electron microscopy (cryo-EM) to an average resolution of 3.3 Å (Figure 3A, Figure S1, and Supplementary Table S1). The resulting reconstruction enabled us to build ∼86% of the Ptx2 model (Figure 3B). Because Ptx2 was purified from the *E. coli* cytoplasm and our cryo-EM structure was determined in the absence of detergents, lipids, or lipid nanodiscs, we conclude that this structure likely represents the soluble conformation that Ptx2 adopts during its export by the H2-T6SS.

**Figure 3:**
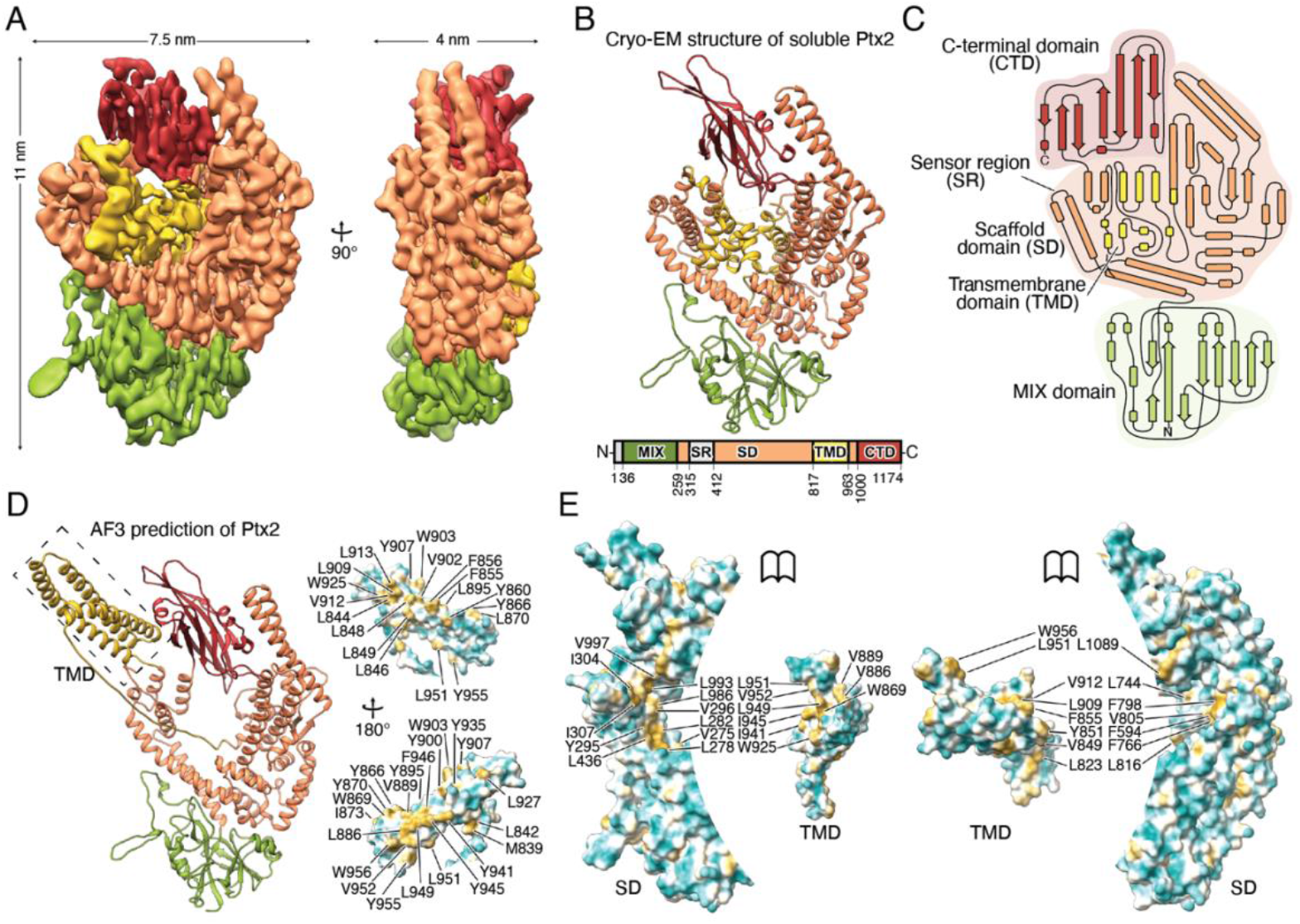
The overall cryo-EM structure of Ptx2. Cryo-EM density (A) and atomic model (B) of Ptx2. A schematic representation of the domain architecture of Ptx2 is provided below the atomic model in (B). C) Schematic representation of the secondary structure of Ptx2. The dotted lines correspond to flexible regions that have not been built. D) AlphaFold 3 prediction of Ptx2 with exposed conformation of the transmembrane domain. SR is hidden for clarity of comparison with (B). E) Shielding of hydrophobic residues (yellow) of the putative transmembrane domain by the hydrophobic interior of the scaffold domain. CTD – C-terminal domain, TMD – putative transmembrane domain, SR – sensor region, SD – scaffold domain, MIX – marker for type six (MIX) domain.

Ptx2 is a monomeric protein that forms an elongated, discoid structure with dimensions of 11 × 7.5 × 4 nm (Figure 3A). Overall, the protein is comprised of several distinct domains (Figure 3B). The N-terminal domain, encompassing residues 36-258, is formed by two antiparallel β-sheets surrounded by seven short α-helices (Figure 3B, C). This domain belongs to the <u>m</u>arker for type s<u>ix</u> (MIX) domain superfamily, which are present in a wide range of structurally and functionally diverse T6SS effectors (Salomon et al., 2014). Despite the extensive study of MIX domain-containing T6SS effectors to date, the structure and precise molecular function of this domain has remained elusive (Dar et al., 2022; Dar et al., 2018; Fridman et al., 2022; Ray et al., 2017; Salomon *et al*., 2014). Because MIX domains are essential for the T6SS-dependent export of effectors that harbor them, it is postulated that this domain serves as an N-terminal secretion signal that enables effector recognition by the T6SS (Fridman *et al*., 2022; Velázquez *et al*., 2025). Consistent with its previously described modular nature, the MIX domain of Ptx2 does not participate in extensive inter-domain interactions like the other domains of the protein. This observation suggests that the MIX domain likely does not directly contribute to the toxic activity of Ptx2 but instead functions in effector recognition and export by the T6SS.

Besides the MIX domain, the majority of Ptx2 forms an almost entirely α-helical domain comprising residues 259-816 and 963-999, which we have termed the scaffold domain. Apart from a flexible region, which we later term the <u>s</u>ensor region (SR; amino acids 315-411), the scaffold domain is well-structured and is formed by a large bundle of 22 α-helices and one 2-stranded antiparallel β-sheet that are tightly packed against one another (Figure 3B, C). Overall, the scaffold domain forms a claw-like structure that entirely encloses a putative trans<u>m</u>embrane <u>d</u>omain (TMD; amino acids 817-962) on three sides and abuts the C-terminal domain of the protein (Figure 3A, B, C). This C-terminal domain adopts an immunoglobulin-like fold consisting of two anti-parallel β-sheets sandwiched against one another (Figure 3C). We queried full-length Ptx2 and each of its individual domains against the DALI structural similarity server and found that they do not resemble proteins of known function. The highest scoring result from this search was the immunoglobulin-like domain of Rab geranylgeranyl transferase (Z-score 5.2, RMSD 9.2 Å over 26 aligned residues), which shares limited topological similarity with the C-terminal immunoglobulin-like domain of Ptx2 (Zhang et al., 2000). We therefore conclude that the N-terminal MIX domain, scaffold, and transmembrane domains of Ptx2 adopt unique protein folds.

We next predicted the structure of Ptx2 using AlphaFold3 and compared this prediction to our cryo-EM model (Abramson et al., 2024). Encouragingly, AlphaFold3 correctly predicts the N-terminal MIX domain, scaffold domain, and C-terminal domain with an overall root mean square deviation (RMSD) of 4 Å. (Figure 3D). However, the position of the putative TMD (residues 817-962) differs dramatically between our cryo-EM structure and the AlphaFold3 prediction (Figure 3D). In our experimental structure, this domain forms 10 short α-helices buried within the palm of the claw-like scaffold domain (Figure 3B, C, E). By contrast, AlphaFold3 predicts that the TMD is formed by three long α-helices arranged in a coiled-coil that protrudes outwards from the plane of the scaffold domain (Figure 3D). This extended coiled-coil conformation exposes numerous hydrophobic residues to the aqueous milieu that are shielded by the scaffold domain in our cryo-EM structure, suggesting that these residues in the AlphaFold3 model are adopting a membrane insertion-competent state (Figure 3D, E). Quantification of this substantial reorganization of the TMD between our cryo-EM structure and the AlphaFold3 model indicates that it rotates by approximately 150° and extends ∼100 Å outwards from its original buried position whereas the conformation of the other three domains of Ptx2 remains similar between these two models. These stark differences in the positioning of the TMD suggest that that the AlphaFold3 model may more closely represent the active, membrane-inserted state of Ptx2 whereas our cryo-EM structure represents Ptx2 in its soluble, pre-membrane insertion conformation.

To experimentally probe the hypothesis that the toxic activity of Ptx2 resides within its putative TMD, we introduced single amino acid mutations into this region of the protein and tested the capacity of these mutants to inhibit the growth of *E. coli* when targeted to the periplasm (Figure 4A). We introduced several mutations into the putative TMD that substitute small, uncharged residues with residues containing bulky or charged side chains, reasoning that these substitutions would be most likely to either disrupt conductance of ions through the Ptx2 pore or destabilize the membrane-inserted conformation of Ptx2. Indeed, expression of these variants in the periplasm of *E. coli* results in reduced growth inhibitory activity and thus provides support for our hypothesis that this putative transmembrane domain region is responsible for the lethal function of this protein (Figure 4B).

**Figure 4:**
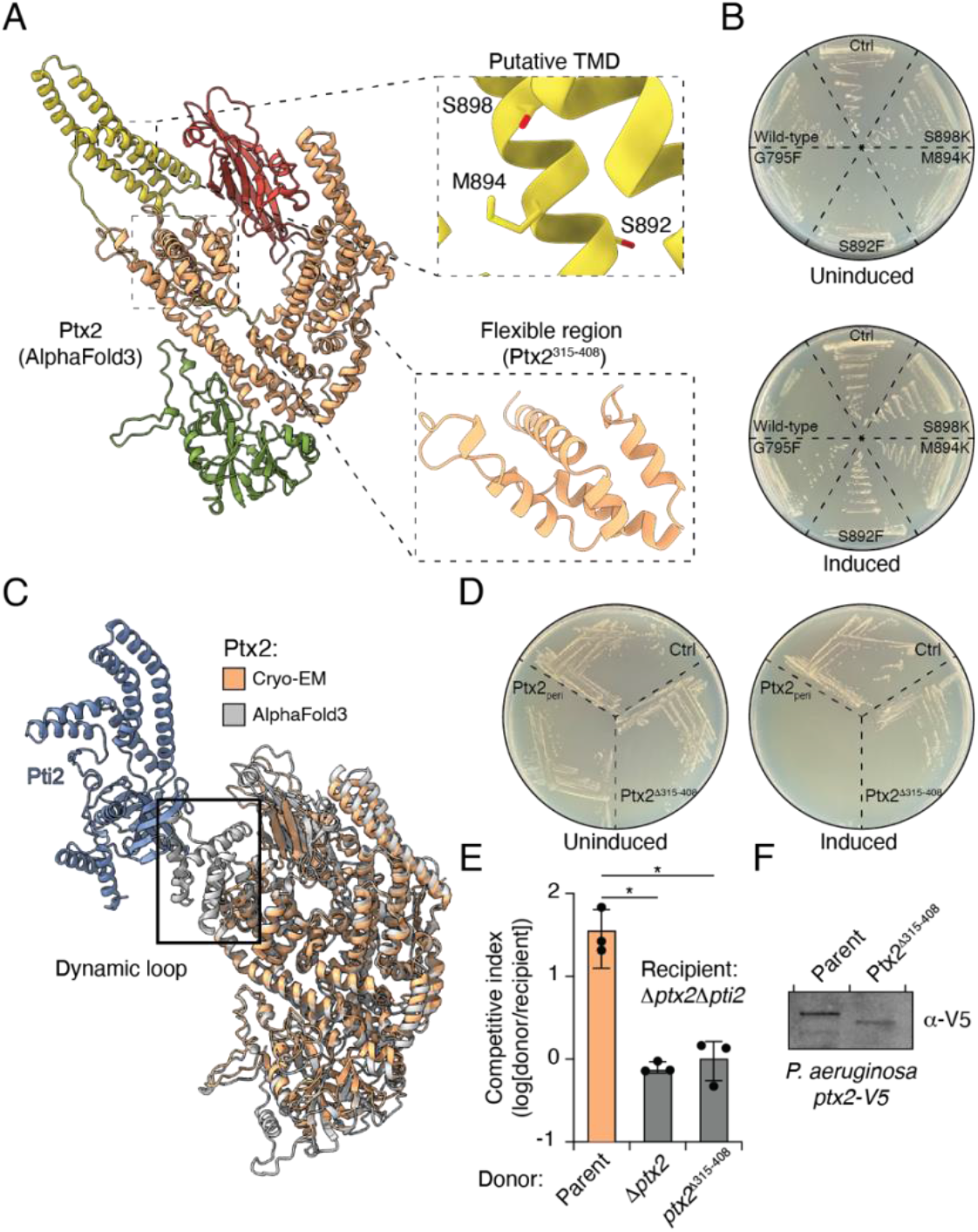
The putative TMD and a dynamic α-helical sensor region of the scaffold domain are required for Ptx2 toxicity. A) The AlphaFold3-predicted structure of Ptx2, which likely represents the active state of this toxin. The inserts highlight residues in the TMD domain required for toxicity (top) and the flexible sensor region predicted to interact with Pti2 (bottom. B, C) AlphaFold3 model of the complex formed by Ptx2 and Pti2, overlaid with the cryo-EM structure of Ptx2 in the soluble conformation. The dynamic sensor region predicted to mediate the Ptx2-Pti2 interaction is highlighted. D) *E. coli* strains harbouring the indicated expression vectors grown on solid media under inducing or non-inducing conditions. E) Outcome of intraspecific growth competitions between the indicated *P. aeruginosa* PA14 donor and recipient strains. The competitive index is calculated as a change (final/initial) in the donor-to-recipient ratio. Error bars represent mean +/-SEM, n=3 biological replicates. Asterisks indicate statistically significant differences between the indicated donor and recipient strain (*p*<0.05). Differences between groups were calculated with an ordinary one-way ANOVA with multiple comparisons to the parent (control) strain. F) Western blot analysis of expression of the indicated *ptx2* alleles encoded at the native chromosomal locus in *P. aeruginosa*.

### A dynamic sensor region in Ptx2 is required for its toxic activity

Because T6SS immunity proteins typically function by physically interacting with domains in their cognate toxins that are essential for toxin function, we reasoned that a structural view of the complex formed between Ptx2 and Pti2 might shed further light on regions of Ptx2 that are responsible for its toxic activity. Despite extensive efforts, we were unable to co-purify a stable complex of Ptx2 and Pti2 for structural studies. We therefore attempted to predict the structure of Ptx2 in complex with Pti2 using AlphaFold3 (Abramson *et al*., 2024). Remarkably, the predicted structure of Ptx2 within this complex strongly resembles our Ptx2 cryo-EM structure with an overall RMSD of 3 Å (Figure 4C, S2A). Unlike the AlphaFold3 prediction of Ptx2 alone where the TMD adopts an extended, solvent-exposed conformation, AlphaFold3 predicts that the TMD of Ptx2 within the Ptx2-Pti2 complex adopts a retracted conformation buried within the scaffold domain (Figure 4C). Furthermore, residues 316-411 of Ptx2 form an α-helical bundle that interacts with a curved four-stranded antiparallel β-sheet in Pti2 (Figure S2B-C). Interestingly, these predicted Pti2-binding α-helices were not resolved in our cryo-EM density, indicating that they are dynamic when Ptx2 is in its soluble state and not inhibited by Pti2. This observation, together with our AlphaFold3 model of the Ptx2-Pti2 complex, led us to hypothesize that this dynamic region is likely responsible for initiating the interaction between Ptx2 and bacterial membranes prior to TMD insertion.

To test our hypothesis that this dynamic region is responsible for Ptx2 toxicity, we engineered a variant of Ptx2 in which we replaced this region with a glycine-serine linker (Ptx2^Δ315-408^) and tested the capacity of this variant to inhibit *E. coli* growth when targeted to the periplasm (Figure 4C). Consistent with our hypothesis that the dynamic region of Ptx2 is responsible for membrane binding, Ptx2^Δ315-408^ did not inhibit *E. coli* growth in this heterologous expression system (Figure 4D). Additionally, we introduced the *ptx2*^Δ315-408^ allele onto the *P. aeruginosa* chromosome at the native *ptx2* locus and tested the competitive fitness of this strain against our Ptx2-susceptible recipient. In line with our findings in *E. coli, P. aeruginosa* harbouring the *ptx2*^Δ315-408^ allele displays no fitness advantage against a Ptx2-susceptible competitor strain, indicating that residues 315-408 are essential for Ptx2 function in vivo (Figure 4E). To confirm that this loss of function is not a consequence of substantially diminished protein expression, we fused a V5 epitope tag to the C-terminus of both wild-type Ptx2 and Ptx2^Δ315-408^ encoded at the native chromosomal locus in *P. aeruginosa* and analyzed expression of these proteins by Western blotting, which confirmed that both Ptx2 variants are expressed in vivo (Figure 4F). We therefore conclude that the dynamic region of the scaffold domain spanning residues 315-408 is essential for the lethal activity of Ptx2.

### The N-terminal MIX domain of Ptx2 binds to its cognate adaptor protein Tap6

Having identified structural features of Ptx2 that contribute to its toxic function, we next sought to determine the function of the MIX domain at the N-terminus of this protein. Because MIX domains have previously been implicated in T6SS effector export, we hypothesized that this domain functions in the recognition of Ptx2 by other components of the T6SS (Fridman *et al*., 2022). Export of Ptx2 relies on a type VI <u>a</u>daptor <u>p</u>rotein (Tap), Tap6, which binds to this effector with high specificity and links it to the secreted T6SS spike protein, VgrG6 (Colautti *et al*., 2024). Homologs of Tap6 co-occur with T6SS effectors that contain MIX domains, further suggesting that Tap6 binds to the N-terminal MIX domain of Ptx2 and recruits it for export by the T6SS (Colautti *et al*., 2024). To better understand the interactions between these proteins, we first sought to determine the structure of the Tap6-Ptx2 complex using cryo-EM. Unfortunately, the small number of particles forming the complex and the apparent flexibility of the interaction prevented us from obtaining a 3D reconstruction. However, we observed additional density near the N-terminal MIX domain of Ptx2 at the level of 2D class averages, which is consistent with our hypothesis that Tap6 binds to the MIX domain.

To directly assess binding between the MIX domain of Ptx2 and Tap6, we next conducted a co-purification experiment between Ptx2’s MIX domain and Tap6. Although the full-length MIX domain was found to be insoluble (residues 36-259), we were able to express and purify a soluble fragment of this domain encompassing residues 32-163. To determine if this MIX domain fragment binds Tap6, we co-expressed a His_6_-tagged version of the protein (Ptx2^32-163^) with untagged Tap6 in *E. coli* and subjected this mixture to co-purification using nickel affinity and size exclusion chromatography. In line with our electron microscopy findings, both full-length Ptx2 and its isolated MIX domain form a stable complex with Tap6 (Figure 5B). This observation confirms our electron microscopy results and leads us to conclude that Tap6 recognizes Ptx2 through its N-terminal MIX domain.

**Figure 5:**
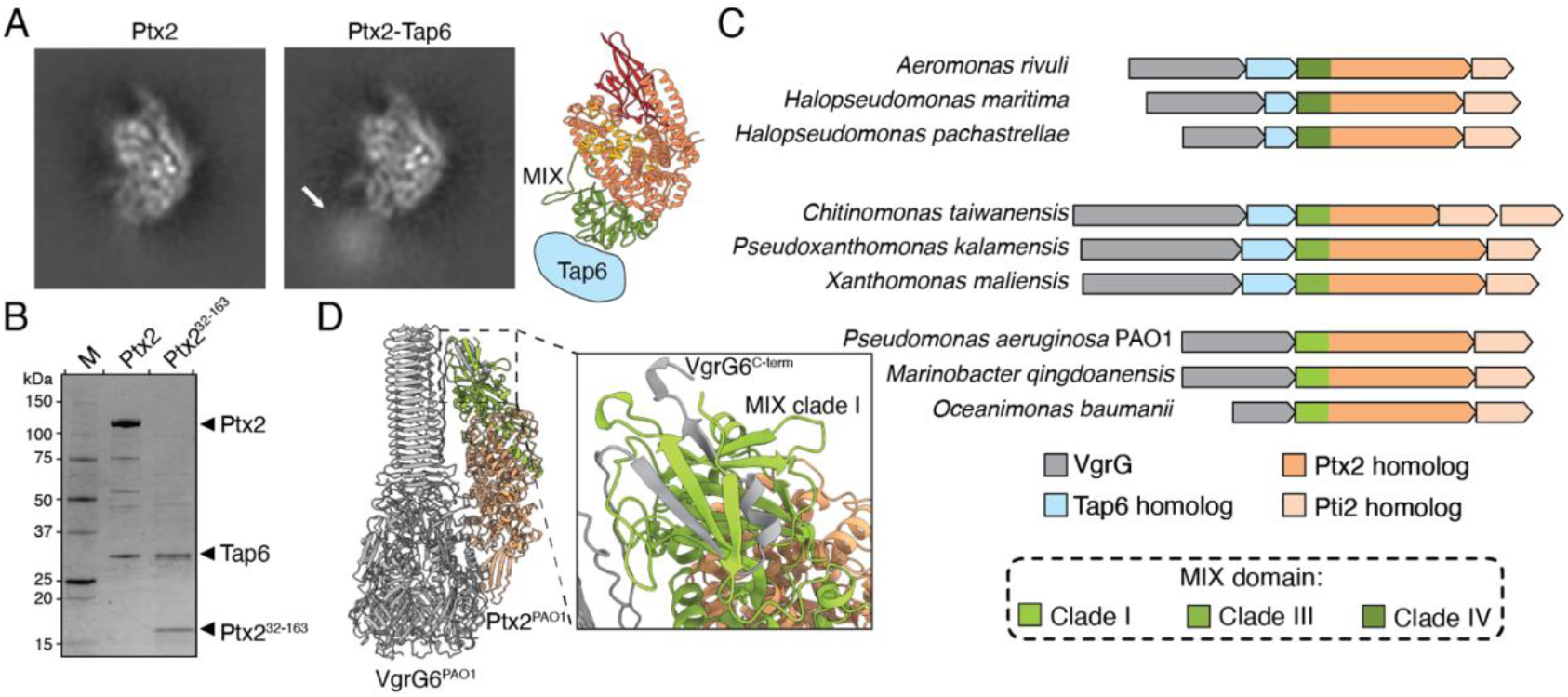
Tap6 binds to the N-terminal MIX domain of Ptx2. A) 2D class averages of Ptx2 alone (left) or Ptx2 in complex with Tap6. Tap6 density is indicated by the white arrow. Cartoon representations of Ptx2 with the domains coloured according to the schematic in figure 3B are presented on the right. B) SDS-PAGE analysis of co-purification between Ptx2 and the MIX domain (Ptx2^32-163^) and Tap6. C) Schematics of Ptx2 homologs that are encoded adjacent to a Tap protein and harbour a MIX domain belonging to clade III or IV of the MIX superfamily and Ptx2 homologs that contain a MIX domain belonging to clade I of the superfamily and are not encoded alongside a Tap protein. D) AlphaFold3 model of the complex formed by VgrG6 and the Ptx2 homolog encoded by *P. aeruginosa* PAO1. The interaction between the clade I MIX domain of Ptx2^PAO1^ and VgrG6^PAO1^ is highlighted in the insert.

Lastly, we aimed to determine whether the interaction between Tap6 and MIX domains represents a widespread mechanism of effector recruitment to the T6SS. To this end, we identified homologs of Ptx2 encoded across the phylum Pseudomonadota and examined the genomic contexts in which these proteins are encoded. The majority of Ptx2 homologs we identified are encoded downstream of a Tap family protein and contain an N-terminal domain belonging to the MIX superfamily, consistent with our finding that Tap binds to the MIX domain to recruit these effectors for T6SS export (Figure 5C, S3A). Interestingly, a subset of Ptx2 homologs are not encoded downstream of a Tap protein, suggesting that these proteins are recruited to the T6SS by a Tap-independent mechanism. To better understand this alternative mechanism of recruitment, we examined the MIX domains present at the N-terminus of these Ptx2 homologs. The MIX superfamily comprises five distinct clades, each defined by the presence of unique conserved sequence motifs (Salomon *et al*., 2014). This diversity suggests that MIX domains belonging to distinct clades may perform related but distinct functions, such as binding to different components of the T6SS apparatus. Consistent with this idea, examination of the domain architecture of the Tap-independent Ptx2 homologs revealed that they exclusively contain clade 1 MIX domains whereas homologs that co-occur with Tap proteins, including Ptx2, contain MIX domains belonging to clades III and IV (Figure 5C, S3B). This strict correlation between MIX domain clade and the presence or absence of a Tap-encoding gene suggests that MIX domains belonging to distinct clades interact with unique binding partners to recruit T6SS effectors for secretion. To explore this possibility, we used AlphaFold3 to predict protein-protein interactions that likely occur between Tap-independent Ptx2 homologs and their associated VgrG spike proteins. Remarkably, AlphaFold3 confidently predicts that the two β-strands found at the extreme C-terminus of a Tap-independent VgrG6 homolog, VgrG6^PAO1^, intercalate with a β-sheet in the clade I MIX domain of its cognate effector, Ptx2^PAO1^ (Figure 5D). This finding supports our hypothesis that the presence of a clade I MIX domain enables Ptx2 homologs to directly interact with their associated VgrG spike protein, thus obviating the need for a Tap adaptor protein. Together, these findings indicate that MIX domains function as N-terminal recognition domains that enable binding of effectors to conserved components of the T6SS spike and implies that this mechanism of VasX family effector recruitment is likely widespread in Pseudomonadota.

## Discussion

Several different families of membrane-depolarizing antibacterial effectors are known to transit the T6SS in diverse bacterial species (Gonzalez-Magana *et al*., 2022; LaCourse *et al*., 2018; Mariano *et al*., 2019; Miyata *et al*., 2013; Velázquez *et al*., 2025). However, the molecular structure of any of these proteins had remained unknown. Here, we use cryogenic electron microscopy to determine the structure of Ptx2, a membrane-depolarizing antibacterial T6SS effector belonging to the VasX protein family. In addition to revealing that Ptx2 does not bear structural similarity to other proteins of known function, our structure enabled the identification of key domains and residues that are responsible for its antibacterial activity. Furthermore, we demonstrate that Ptx2 is recruited to the T6SS apparatus through its N-terminal MIX domain, which provides insight into the molecular role these domains play in the export of T6SS effectors that harbor them.

Proteins belonging to the VasX family have long been known to function as membrane-depolarizing T6SS effectors (Miyata *et al*., 2011; Miyata *et al*., 2013; Russell *et al*., 2012; Velázquez *et al*., 2025). VasX, the first protein in this family to be characterized, was originally identified as a virulence factor required for *V. cholerae* pathogenesis in the model eukaryotic organism *Dictyostelium discoideum* (Miyata *et al*., 2011). VasX was subsequently demonstrated to act as an ion channel that dissipates the proton motive force (Miyata *et al*., 2013). This mechanism of action explained the surprising finding that VasX acts both as an antibacterial T6SS effector and as a T6SS-delivered virulence factor because transmembrane ion gradients play an important role in the cellular physiology of both bacterial and eukaryotic cells (Miyata *et al*., 2013). Like VasX, the *P. putida* T6SS effector Tke5 functions as a membrane-depolarizing antibacterial toxin and was further found to form cation-selective channels in the bacterial cytoplasmic membrane (Velázquez *et al*., 2025). The similarity between VasX and Tke5 extends to the finding that like VasX, Tke5 can kill a wide range of susceptible bacteria. In addition to these two extensively characterized prototypes and the less well-studied *B. thailandensis* effector BTH_I2691, VasX family T6SS effectors are widespread among the phylum Pseudomonadota, which suggests that these proteins play an important role in interbacterial antagonism and virulence in multiple ecological contexts (Miyata *et al*., 2013; Russell *et al*., 2012; Velázquez *et al*., 2025).

Our cryo-EM structure of Ptx2 offers a molecular view of the VasX protein family and our comparison to an AlphaFold3 generated model of the protein enabled the identification of key residues responsible for its toxicity. In light of these findings, we propose the following model for Ptx2 function, which we anticipate will be consistent across the VasX protein family: in the cytoplasm of the producing cell, a tripartite complex forms between the secreted spike protein VgrG6, the adaptor protein Tap6, and the N-terminal MIX domain of Ptx2 in its soluble conformation (Figure 6). Following secretion into a susceptible recipient, the dynamic sensor region of Ptx2 (residues 315-411) interacts with the cytoplasmic membrane and initiates a conformational change in which the buried TMD unfolds into its extended, membrane-inserted state. This membrane-inserted conformation of Ptx2 conduct ions across the cytoplasmic membrane, thereby dissipating the PMF and inhibiting bacterial growth. If Ptx2 is delivered into a recipient cell that harbours the cognate immunity protein, Pti2, then we expect that this immunity protein will bind to the dynamic α-helices of the Ptx2 scaffold domain, thus preventing the conformational changes required for Ptx2 insertion into the cytoplasmic membrane.

**Figure 6:**
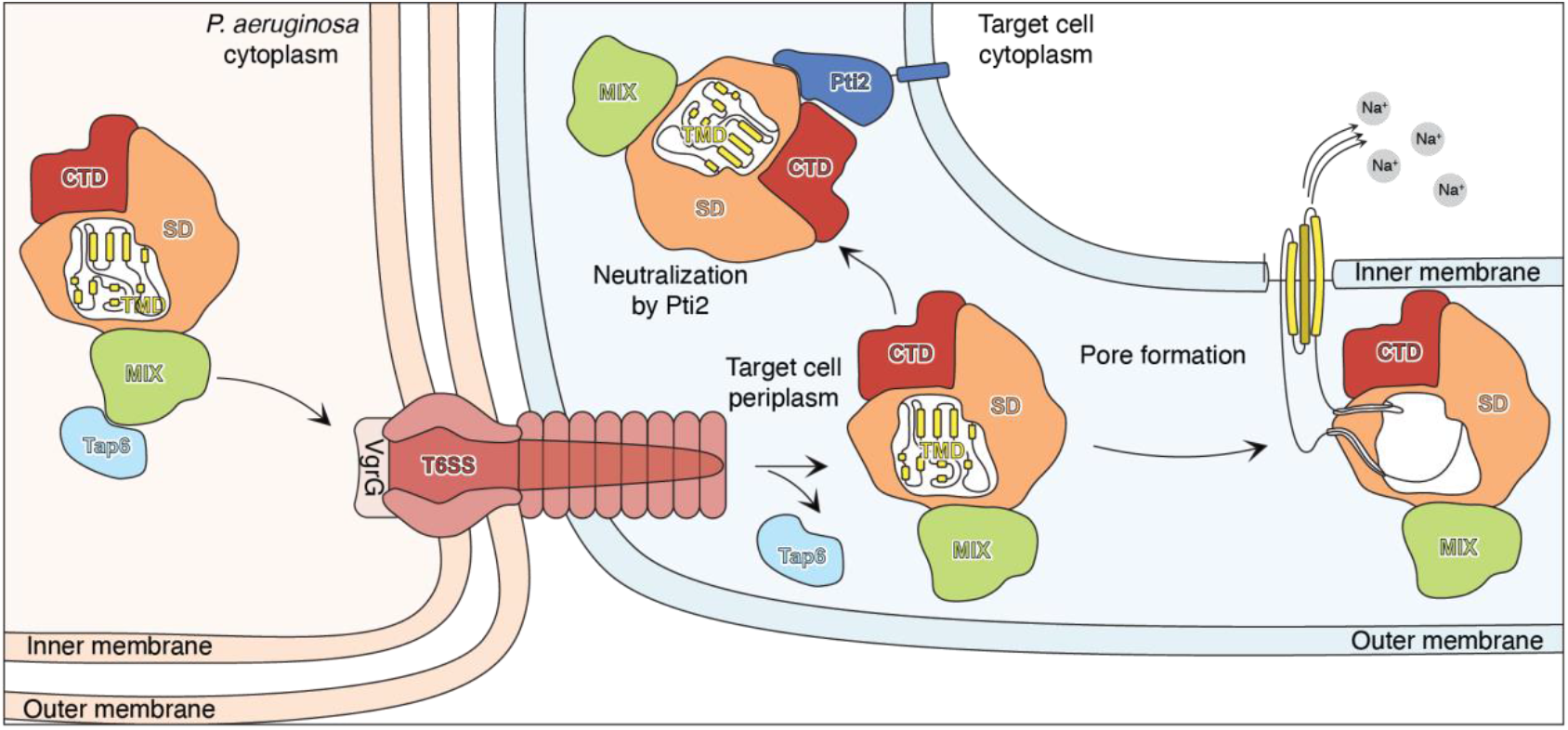
Model of Ptx2 toxicity and delivery by the T6SS. In the cytoplasm of *P. aeruginosa*, Tap6 interacts with the MIX domain of Ptx2 and links it to VgrG6 for T6SS-dependent export into the periplasm of the target cell. Upon delivery into this compartment of a susceptible target cell, Tap6 may dissociate from Ptx2 through an unknown mechanism prior to Ptx2 insertion into the cytoplasmic membrane. Insertion of the Ptx2’s TMD into the membrane results in the formation of an ion-conductive pore, which enables passive influx of sodium ions and thus dissipates the proton motive force. Alternatively, in a target cell expressing Pti2, the membrane-bound immunity protein interacts with Ptx2 and blocks its insertion into the membrane.

Although many steps of this model have been demonstrated by our work and that of others, several aspects of Ptx2 toxicity remain incompletely understood. Perhaps the most important outstanding question concerns the membrane-inserted conformation of this protein. Our mutagenesis analyses suggest that several residues in the putative TMD are essential for the toxicity of Ptx2, although the precise role these residues play in the function of this protein remains to be defined. One possibility is that substitution of these small side chains with lysine or phenylalanine forms a bulky plug within the membrane-inserted ion channel, thus blocking ion conduction through the Ptx2 pore. Alternatively, it is possible that these substitutions form intramolecular interactions with nearby residues in the retracted TMD, thus stabilizing the soluble state of Ptx2 and preventing the conformational changes required to insert into the cytoplasmic membrane. Additionally, it remains unclear whether Ptx2 forms a monomeric ion channel or oligomerizes upon inserting into the membrane as is the case for many other families of membrane-depolarizing antibacterial proteins (Dal Peraro and van der Goot, 2016). However, because prior work on the mechanism of Ptx2 secretion by the T6SS revealed that a single T6SS secretion event can export, at most, three copies of this toxin, it seems unlikely that Ptx2 would function in an oligomeric state greater than a trimer (Colautti *et al*., 2024). Further structural analyses that capture the membrane-inserted state of Ptx2 or other VasX family proteins may shed light on these mechanistic questions. Although we were able to purify Ptx2 from the membrane fraction of *E. coli* and solubilize this protein in detergent micelles, the protein yield was insufficient for structural studies using cryo-EM. More sophisticated approaches, such as preparation of Ptx2 in lipid nanodiscs or liposomes, may therefore be required to prepare samples of sufficient quantity and quality for high-resolution structure determination.

In conclusion, we determined the cryo-EM structure of a membrane-depolarizing antibacterial T6SS effector exported by *P. aeruginosa*. This protein, which belongs to the extensively characterized VasX effector family, adopts a highly unique structure comprising multiple domains that bear little similarity to proteins of known function. Our findings provide molecular insight into the mechanisms by which effectors in this family are recruited to the T6SS for export and how they exert their toxic effects upon delivery to target bacteria.

## Methods

### Bacterial strains and culture conditions

*P. aeruginosa* strains were derived from the reference strain PA14 (Lee *et al*., 2006). *E. coli* strain XL1 Blue (Novagen) was used for plasmid maintenance, SM10 was used for conjugative transfer, and BL21 (DE3) pLysS (Novagen) was used for protein expression. Cultures were grown in lysogeny broth (LB) medium (10 g/L tryptone, 5 g/L yeast extract, 10 g/L NaCl) at 37°C shaking at 220 RPM. Cultures were supplemented with 50 μg/mL kanamycin, 100 μg/mL ampicillin, 25 μg/mL chloramphenicol, 15 μg/mL (*E. coli*) or 30 μg/mL (*P. aeruginosa*) gentamicin, 25 μg/mL irgasan (*P. aeruginosa*), 0.1% (w/v) L-arabinose, or 1 mM β-D-thiogalactopyranoside (IPTG), as appropriate. A detailed list of strains used in this study can be found in table S1.

### DNA manipulation, plasmid construction, and mutant strain generation

Plasmids were constructed using standard restriction enzyme cloning methods (Green et al., 2012). All primers were synthesized by Integrated DNA Technologies (IDT). Phusion polymerase, restriction endonucleases, and T4 DNA ligase was obtained from New England Biolabs (NEB). Sanger sequencing was performed by The Center for Applied Genomics (TCAG) at the Hospital for Sick Children (Toronto, Ontario) for plasmids containing cloned inserts <1000 bp. Plasmids containing cloned inserts >1000 bp were sequenced by Plasmidsaurus using Oxford Nanopore technology. A complete list of plasmids used in this study can be found in table S2.

Chromosomal mutations were generated in *P. aeruginosa* using the double allelic exchange method as previously described (Hmelo et al., 2015). Approximately 500 bp flanks upstream and downstream of the region to be mutated were amplified by PCR and spliced together by overlap extension PCR, with the desired mutation introduced between these flanks by overlap extension. The resulting amplicon was ligated into the allelic exchange vector pEXG2, transformed into the conjugative *E. coli* strain SM10, and introduced to *P. aeruginosa* by conjugation. Merodiploids were selected at 37°C on LB agar containing 30 μg/mL gentamicin and 25 μg/mL irgasan and subsequently streaked on LB lacking NaCl containing 5% (w/v) sucrose at 30°C. Strains that grow on sucrose and are gentamicin sensitive were screened by colony PCR to specifically amplify the mutated allele. Chromosomal mutations were verified by PCR amplification of the region containing the mutation and Sanger sequencing of the resulting amplicon.

### Bacterial competition assays

Intraspecific competition assays were performed as previously described (Colautti *et al*., 2024). The indicated donor and recipient strains were grown overnight in LB medium at 37°C with shaking at 220 RPM. Recipient strains harbour a constitutively expressed *lacZ* reporter gene integrated at a neutral chromosomal locus. Stationary phase cultures were normalized to an OD_600_ of 1.0 and donor and recipient cultures were combined to an initial donor/recipient ratio of 5:1. To quantify initial colony forming units of both donor and recipient strains, 10-fold serial dilutions of this mixture were plated on LB containing 40 μg/mL 5-bromo-4-chloro-3-indolyl-β-D-galactopyranoside (X-gal) and incubated at 37°C for 18 hours. Simultaneously, 10 μL of the initial donor/recipient mixture was spotted onto a nitrocellulose membrane overlaid onto LB containing 3% agar, which were incubated upright for 20 hours at 25°C. Where indicated, LB 3% (w/v) agar contained 150 mM KCl instead of NaCl, or lacked NaCl and contained 300 mM sucrose. The pH of LB 3% (w/v) agar was adjusted to the indicated pH using sodium phosphate. Final CFUs were quantified by resuspending the resulting colonies in 1 mL LB medium and plating 10-fold serial dilutions of this suspension on LB containing 40 μg/mL X-gal. Competitive index is calculated as the ratio of donor/recipient at the final timepoint divided by the initial donor/recipient ratio.

Monoculture competition experiments were performed as previously described (Russell *et al*., 2011). The indicated strains were grown overnight in LB medium at 37°C with shaking at 220 RPM. Stationary phase cultures were normalized to an OD_600_ of 1.0, 10 μL were spotted onto nitrocellulose membranes overlaid on LB medium containing 3% (w/v) agar and incubated at 25°C for 20 hours. To quantify growth on this condition, the resulting colonies were resuspended in 1 mL LB medium and 10-fold serial dilutions of this suspension were plated on LB medium containing 1.5% (w/v) agar and grown at 37°C for 18 hours. Results are reported as CFU/mL.

### P. aeruginosa growth curves

*P. aeruginosa* strains were grown overnight at 37°C shaking at 220 RPM in 2 mL LB. Cultures were diluted 200-fold in LB, and 200 μL of this diluted culture was allowed to grow for 18 hours in a 96-well microtiter plate. Growth curves were performed at 25°C, a temperature that induces expression of T6SS genes (Allsopp *et al*., 2017), in a BioTek Epoch plate reader measuring OD_600_ every 30 minutes.

### Protein expression and purification

*E. coli* BL21 (DE3) pLysS (Novagen) harbouring the indicated expression vector was grown at 37°C shaking at 220 RPM in LB medium supplemented with appropriate antibiotics to an OD_600_ of 0.6. Cultures were cooled to 18°C and protein expression was induced by the addition of IPTG to a final concentration of 1 mM. Proteins were expressed at 18°C for 18-20 hours before cells were collected by centrifugation at 6000x*g* for 20 minutes at 4°C. Cells were resuspended in lysis buffer (50 mM HEPES NaOH pH 7.5, 300 mM NaCl) and lysed by sonication. Lysates were clarified by centrifugation at 35 000x*g* for 45 minutes at 4°C before being loaded onto a 1 mL Ni-NTA agarose column pre-equilibrated with 5 mL wash buffer (50 mM HEPES NaOH pH 7.5, 300 mM NaCl, 10 mM imidazole). Columns were washed 3x with 20 mL wash buffer before His_6_-tagged proteins were eluted by the addition of 4 mL elution buffer (50 mM HEPES NaOH pH 7.5, 300 mM NaCl, 400 mM imidazole). Eluted proteins were further purified by gel filtration on a HiLoad 16/600 Superdex 200 size exclusion chromatography column (GE healthcare) on an AKTA system. Gel filtration was performed using SEC buffer (20 mM HEPES NaOH pH 7.5, 150 mM NaCl). The purity of each protein was determined using SDS-PAGE followed by staining with Coomassie brilliant blue R250. Proteins were concentrated to 10 mg/mL, snap frozen in liquid nitrogen, and stored at -80°C for further use.

### Cryogenic electron microscopy

3 µl of Ptx2 Ptx2-Tap6 complex at 0.4 mg/ml was applied onto a freshly glow-discharged copper R1.2/1.3 300 mesh grid (Quantifoil), blotted for 3 s on both sides with blotting force 0 and plunge-frozen in liquid ethane-propane mixture using the Vitrobot Mark IV system (Thermo Fisher Scientific) at 13 °C and 100% humidity. The cryo-EM datasets were collected using a Krios transmission electron microscope (Thermo Fisher Scientific) equipped with an XFEG at 300 kV using the automated data-collection software EPU. One image per hole with defocus range of -0.5 - -2.5 µm was collected with a K3 detector (Gatan) operated in super-resolution mode. Image stacks with 50 frames were collected with a total exposure time of 1.75 sec and a total dose of 50 e-/Å^2^. 6997 movies of Ptx2 and 1807 movies of Ptx2-Tap6 were used for data processing. After motion correction and CTF estimation in cryoSPARC (Punjani et al., 2017), we picked particles using blob picker setting the diameter range to 90-120 Å and extracted them with the box size 196 px. After 2D classification, good classes were used as templates for picking from the same micrographs, particles were also extracted with box size 196, and 2D classified. Good particles from both 2D classifications were pulled together, and duplicates with distance less than 100 Å were removed. The remaining particles were reextracted with box size 256, classified once again in 2D, and selected particles were used for ab initio 3D reconstruction, followed by a non-uniform refinement. Despite seeing Tap6 density in some 2D classes in the Ptx2-Tap6 dataset (Figure 5A, S1H), we were unable to identify Tap6 density in 3D reconstruction due to the low abundance of the complex and potential flexibility at the interaction site. Multiple 3D classifications did not allow us to separate Ptx2-Tap6 particles from Ptx2. Therefore, we continued to process only the Ptx2 dataset. Initial Ptx2 reconstruction showed clear anisotropy due to preferred position of particles on ice. To compensate it, we removed ∼40% of particles from overpopulated views and repeated a non-uniform 3D refinement with the remaining particles. Using this 3D reconstruction, we performed global and local CTF refinement, and, finally, performed the last 3D refinement in Relion 5 with Blush regularization (Kimanius et al., 2024). The local resolution-filtered map was used for model building. To this end, we predicted the structure of Ptx2 in AlphaFold 3 (Abramson *et al*., 2024), performed rigid body fit in Chimera (Pettersen et al., 2004), and refined the structure in Isolde (Croll, 2018), in ChimeraX (Navarro et al., 2024), and Phenix (Liebschner et al., 2019).

### E. coli toxicity assays

*E. coli* XL1 Blue (Novagen) strain harbouring the indicated pBAD33-derived vectors were streaked on LB 1.5% (w/v) agar containing 25 μg/mL chloramphenicol and grown at 37°C for 18 hours. To minimize analysis of spontaneous suppressors that arise in these *E. coli* strains, a single colony of each strain was streaked on LB 1.5% (w/v) agar containing 25 μg/mL chloramphenicol and either containing or lacking 0.1% L-arabinose. The resulting plates were incubated at 37°C for 18 hours before being photographed.

### SDS-PAGE and Western Blotting

Protein samples were diluted in 4x Laemmli SDS-PAGE loading buffer and boiled at 95°C for 10 minutes. Precipitated material was pelleted by centrifugation at 21 000x*g* for 1 minute and equal volumes of supernatant were loaded onto a 12% polyacrylamide gel. Gels were run at 95 V for 12 minutes, followed by 40 minutes at 195 V. Total protein was wet transferred to a nitrocellulose membrane at 103 V for 33 minutes using a Mini Trans-Blot electrophoretic transfer system (BioRad). Membranes were blocked with Tris-buffered saline containing 0.05% Tween-20 (TBS-T) containing 0.5% (w/v) non-fat milk. Primary antibodies were added at 1:5000 dilution and incubated at room temperature for 1 hour, gently shaking (60 RPM) at room temperature. The blot was washed 3x for 5 minutes each with TBS-T, horseradish peroxidase-conjugated anti-rabbit secondary antibody (New England Biolabs) was added at a 1:5000 dilution. The blot was incubated with secondary antibody for 45 minutes at room temperature shaking at 60 RPM before being washed 3x for 5 minutes each with TBS-T. The blot was developed using Clarity Max ECL substrate (BioRad) and visualized using a chemidoc instrument (BioRad).

### Bioinformatic identification of Ptx2 homologs

Ptx2 homologs were identified using position-specific iterative basic local alignment (PSI-BLAST) (Altschul et al., 1997). The scaffold and putative transmembrane domain of Ptx2 was queried against the RefSeq-select non-redundant protein database using an expect value cutoff of 0.0001. To reduce the likelihood of identifying false positive hits, results were restricted to proteins encoded by genomes belonging to the phylum Pseudomonadota. After 5 iterations of PSI-BLAST, the gene neighbourhoods of the top 500 homologous sequences were analyzed using FlaGs (Saha et al., 2021). We subsequently identified conserved domains within these Ptx2 homologs using the NCBI Conserved Domain Database (Wang et al., 2023).

## Acknowledgements

We thank Michiel Punter for maintaining the computing cluster, and members of the Whitney and Belyy labs for helpful discussions. This work benefited from access to the Netherlands Centre for Electron Nanoscopy (NeCEN) at Leiden University, an Instruct-ERIC centre with assistance from Deivanayagabarathy Vinayagam. Access to NeCEN was partially funded by the Netherlands Electron Microscopy Infrastructure (NEMI), project number 184.034.014 of the National Roadmap for Large-Scale Research Infrastructure of the Dutch Research Council (NWO). JC is supported by a Canada Graduate Scholarship from the Canadian Institutes of Health Research (CIHR). This work was supported by a Project Grant from CIHR to JCW (PJT-175011). JCW is the Canada Research Chair in Molecular Microbiology and holds an Investigators in the Pathogenesis of Infectious Disease Award from the Burroughs Wellcome fund.

## Notes

### Competing Interest Statement

The authors have declared no competing interest.

