## Supporting Information for "Cryo-EM structure of a type VI secretion system delivered membrane-depolarizing toxin involved in bacterial antagonism"

Supplementary Figures S1-S3
Supplementary Table S1-S3
Supplementary References

Keywords: *Pseudomonas aeruginosa*, type VI secretion systems, membrane-depolarizing toxins, cryo-EM

*To whom correspondence should be addressed: Alexander Belyy or John C. Whitney
Telephone – (+31) 6 3198 3128; (+1) 905-525-9140

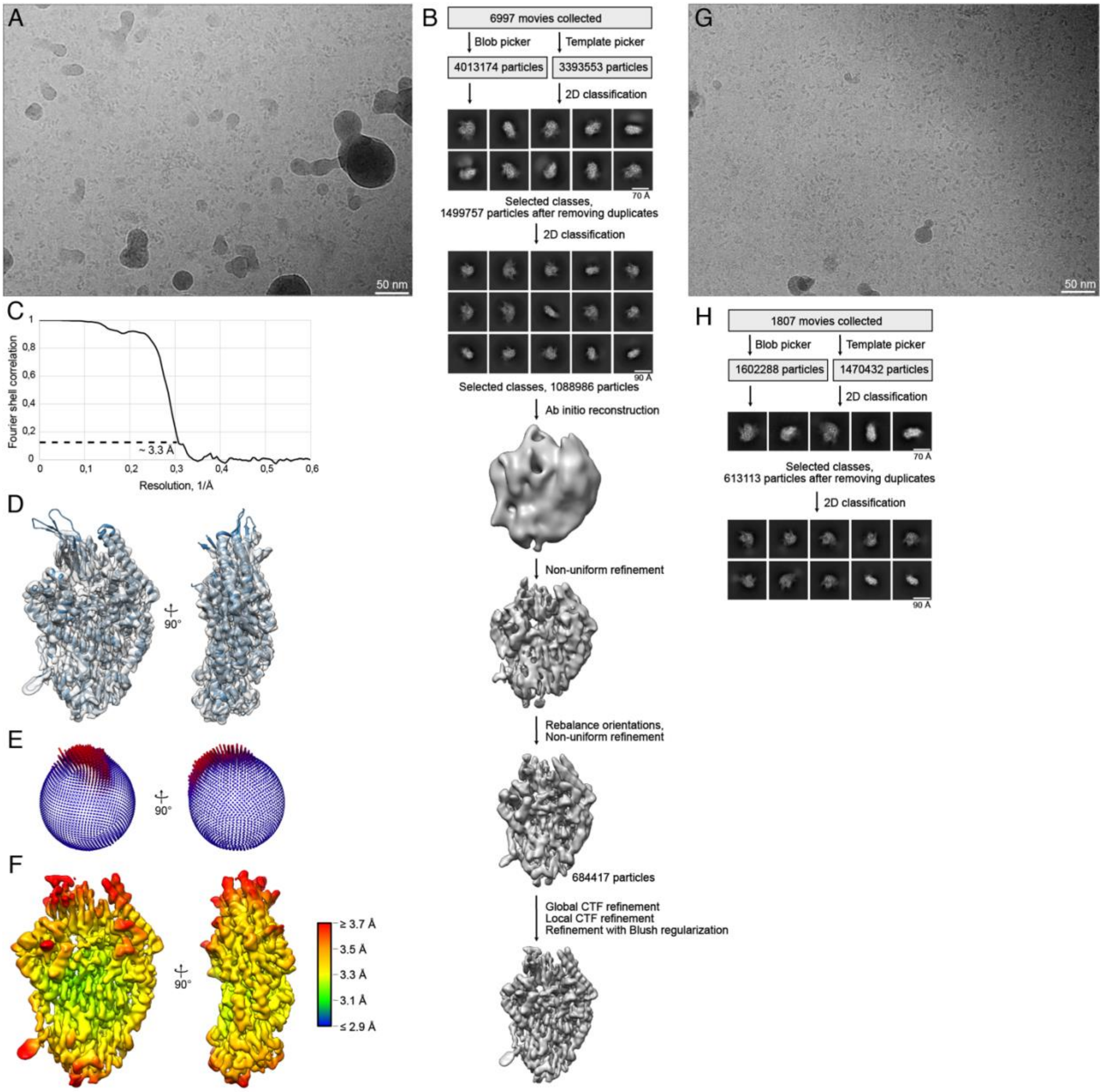


**Supplementary Figure S1: Cryo-EM data processing.** A) Exemplary cryo-EM micrograph of Ptx2. B) Processing overview of the Ptx2 dataset. C) Fourier shell correlation (FSC) curve of the final masked reconstruction. D) Fit of the molecular model into the cryo-EM density. E) Angular distribution of the reconstruction. F) Local resolution gradient of the reconstruction calculated at FSC threshold 0.143. G) Exemplary cryo-EM micrograph of the Ptx2-Tap6 complex. H) Processing of the Ptx2-Tap6 dataset.

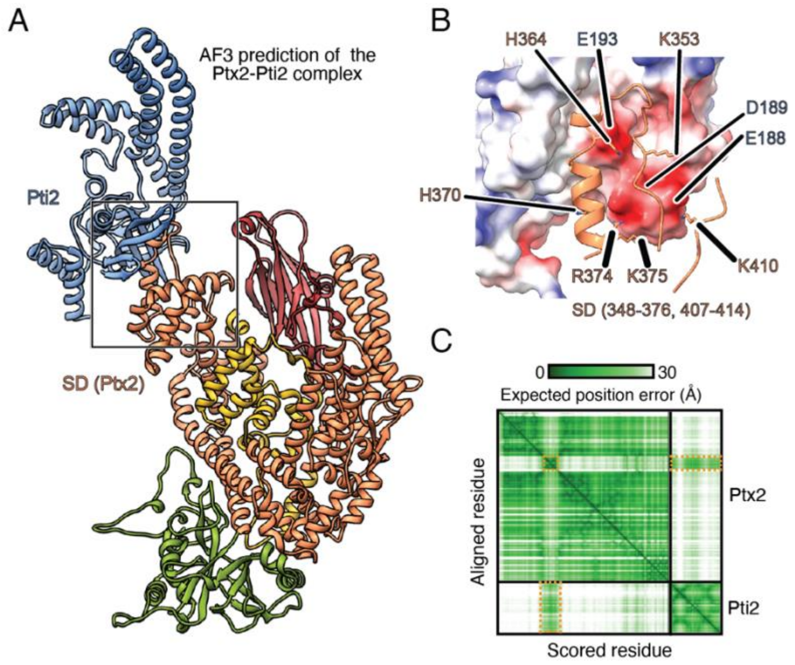


**Supplementary Figure S2: AlphaFold3 predicts that Pti2 interacts with the soluble conformation of Ptx2** A) The AlphaFold3 predicted structure of Ptx2 in complex with Pti2. Domains of Ptx2 are coloured according to the schematic in Figure 3 (B). The predicted interaction between Ptx2 and Pti2 is highlighted in the insert. C) The insert of panel (B) highlighting the predicted interaction interface between Ptx2 and Pti2. Ptx2 is depicted in orange and Pti2 is depicted in surface electrostatic representation with cationic (blue) and anionic (red) surfaces illustrated. D) Predicted aligned error (PAE) plot of the Ptx2/Pti2 complex prediction. Residues corresponding to the predicted interaction interface between Ptx2 and Pti2 are highlighted in orange.

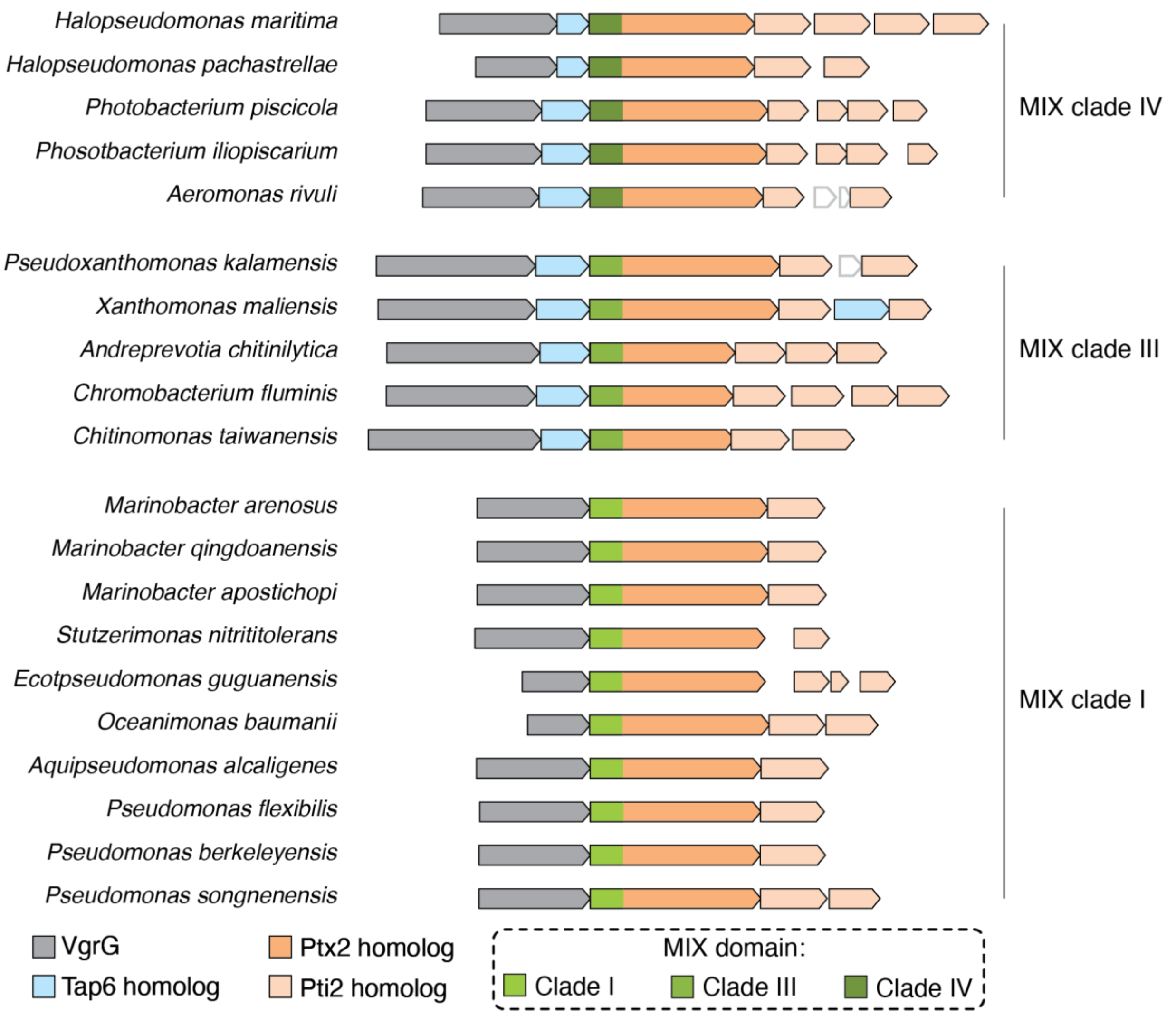


**Supplementary Figure S3: The clade of N-terminal MIX domain correlates with the presence or absence of a co-occurring Tap protein in Ptx2 homologs** Schematic representation of the gene neighbourhoods of Ptx2 homologs encoded by diverse Pseudomonadota. VgrG (gray), Tap (blue), and Ptx2 homologs (orange) are coloured. The clade of MIX domain present in each Ptx2 homolog is indicated.

**Supplementary Table S1:** Cryo-EM data collection, refinement, and validation statistics

| | |
|---|---|
| Microscope | Titan Krios |
| Voltage (kV) | 300 |
| Defocus range (µm) | -0.5 to -2.5 |
| Camera | Gatan K3 Superresolution mode |
| Pixel size (Å) | 0.84 |
| Total electron dose (e/Å$^2$) | 50 |
| Exposure time (s) | 1.75 |
| Frames per movie | 50 |
| Number of movies | 6997 |
| **3D Refinement** | |
| Number of particles | 684417 |
| Final map pixel size | 0.84 |
| Final resolution (Å) | 3.3 |
| **Atomic model statistics** | |
| Non-hydrogen atoms | 7777 |
| Number of chains | 1 |
| Molprobity score | 1.83 |
| Rama distribution Z-score | -2.31 ± 0.24 |
| Clashscore | 7.61 |
| Bond RMSD (Å) | 0.003 |
| Angle RMSD (°) | 0.775 |
| Poor rotamers (%) | 0 |
| Favored rotamers (%) | 99.5 |
| Ramachandran favored (%) | 93.74 |
| Ramachandran allowed (%) | 6.26 |
| Ramachandran outliers (%) | 0 |
| Missing fragments | 1-35, 315-411, 711-718, 891-905, 1009-1020 |

**Supplementary Table S2**: Strains used in this study

| Organism | Genotype | Description | References |
|---|---|---|---|
| *Pseudomonas aeruginosa* PA14 | Wild type | | (Lee et al., 2006) |
| | ΔPA14_52570<br>ΔPA14_20290 | *rsmA amrZ* deletion strain | (Bullen et al., 2022a) |
| | ΔPA14_52570<br>ΔPA14_20290<br>*attB::lacZ* | *rsmA amrZ* deletion, *lacZ* reporter strain | (Ahmad et al., 2019) |
| | ΔPA14_52570<br>ΔPA14_20290<br>ΔPA14_69520<br>ΔPA14_69510<br>*attB::lacZ* | *rsmA amrZ ptx2 pti2* deletion, *lacZ* reporter strain | (Colautti et al., 2024) |
| | ΔPA14_52570<br>ΔPA14_20290<br>ΔPA14_69510 | *rsmA amrZ pti2* deletion strain | This study |
| | ΔPA14_52570<br>ΔPA14_20290<br>ΔPA14_69510<br>ΔPA14_42900 | *rsmA amrZ pti2 tssM2* deletion strain | This study |
| | ΔPA14_52570<br>ΔPA14_20290<br>PA14_69520_Δ315-408 | *rsmA amrZ* deletion, *ptx2*$^{\Delta 315\text{-}408}$ strain | This study |
| | ΔPA14_52570<br>ΔPA14_20290<br>PA14_69520-V5 | *rsmA amrZ* deletion, *ptx2*-V5 strain | This study |
| | ΔPA14_52570<br>ΔPA14_20290<br>PA14_69520_Δ315-408-V5 | *rsmA amrZ* deletion, *ptx2*$^{\Delta 315\text{-}408}$-V5 strain | This study |
| *E. coli* BL21 (DE3) Codon Plus | F- ompT gal dcm lon hsdSB($rB^- mB^-$) λ(DE3) pLysS($cm^R$) | Protein expression strain | Novagen |
| *E. coli* SM10 λpir | *thi thr leu tonA lac Y supE recA*::RP4-2-Tc::Mu | Conjugation strain | BioMedal LifeScience |

| | | | |
|---|---|---|---|
| *E. coli* XL-1 Blue | *recA1 endA1 gyrA96 thi-1 hsdR17 supE44 relA1 lac* [F′ *proAB lacI$^q$* Z*ΔM15* Tn*10* (Tet$^R$)] | Cloning strain | Novagen |

**Supplementary Table S3:** Plasmids used in this study

| Plasmid | Relevant features | Reference |
|---|---|---|
| pEXG2 | Allelic replacement vector containing *sacB,* Gm$^R$ | (Rietsch et al., 2005) |
| pEXG2::ΔPA14_69510 | *pti2* deletion construct | This study |
| pEXG2::PA14_69520_Δ315-408 | *ptx2*$^{\Delta 315\text{-}408}$ allelic exchange construct | This study |
| pEXG2:: ΔPA14_42900 | *tssM2* deletion construct | (Bullen et al., 2022b) |
| pEXG2::PA14_69520-V5 | *ptx2*-V5 construct | (Colautti *et al.*, 2024) |
| pET29b | Expression vector with *lacI*, T7 promoter, C-terminal His$_6$ tag, Kan$^R$ | Novagen |
| pET29b::PA14_69520-His6 | Expression vector for Ptx2 | (Colautti *et al.*, 2024) |
| pET29b::PA14_69520_308-415_GGGSx2-His6 | Expression vector for Ptx2$^{\Delta 315\text{-}408}$. Residues 315-408 in Ptx2 replaced with GGGSGGGS. | This study |
| pETDuet | Co-expression vector with *lacI*, T7 promoter, N-terminal His$_6$ tag in MCS-1, Amp$^R$ | Novagen |

| | | |
|---|---|---|
| pETDuet::His$_6$-PA14_69520::FLAG-PA14_69540 | Co-expression vector for His$_6$-Ptx2 and FLAG-Tap6 | (Colautti *et al.*, 2024) |
| pETDuet::His$_6$-PA14_69520_32-163::FLAG-PA14_69540 | Co-expression vector for His$_6$-Ptx2$^{32\text{-}163}$ and FLAG-Tap6 | This study |
| pBAD33 | Expression vector with *araBAD*, *ara* promoter, ChlorR | (Guzman et al., 1995) |
| pBAD33::PA14_69520 | Ptx2$^{cyto}$ expression vector | This study |
| pBAD33::amiA-PA14_69520 | Ptx2$^{peri}$ expression vector containing the Tat-dependent signal peptide of *E. coli amiA*. | This study |
| pBAD33::amiA-PA14_69520_315-408_GGGSx2 | Ptx2$^{\Delta 315\text{-}408}$ expression vector containing the Tat-dependent signal peptide of *E. coli amiA*. | This study |
| pBAD33::amiA-PA14_69520_G795F | Ptx2$^{G795F}$ expression vector containing the Tat-dependent signal peptide of *E. coli amiA*. | This study |
| pBAD33::amiA-PA14_69520_S892F | Ptx2$^{S982F}$ expression vector containing the Tat-dependent signal peptide of *E. coli amiA*. | This study |
| pBAD33::amiA-PA14_69520_M894K | Ptx2$^{M894K}$ expression vector containing the Tat-dependent signal peptide of *E. coli amiA*. | This study |

| | | |
|---|---|---|
| pBAD33::amiA-PA14_69520_S898K | Ptx2$^{S898K}$ expression vector containing the Tat-dependent signal peptide of *E. coli amiA*. | This study |
